# Novel co-culture strategies of tumor organoids with autologous T-cells reveal clinically relevant combinations of immune-checkpoint and targeted therapies

**DOI:** 10.1101/2023.07.05.546622

**Authors:** Enrique Podaza, Jared Capuano, Majd Al Assaad, Hui-Hsuan Kuo, Geoffrey Markowitz, Adriana Irizarry, Hiranmayi Ravichandran, Sarah Ackermann, Troy Kane, Jyothi Manohar, Michael Sigouros, Jenna Moyer, Bhavneet Bhinder, Pooja Chandra, Murtaza Malbari, Karsten Boehnke, Juan Miguel Mosquera, Vivek Mittal, Andrea Sboner, Hamza Gokozan, Nasser Altorki, Olivier Elemento, M. Laura Martin

## Abstract

Patient derived tumor organoids (PDTOs) have become relevant pre-clinical models for therapeutic modelling since they highly recapitulate patients’ response to treatment. Nevertheless, their value for immunotherapy modelling has not been fully explored. We developed a tumor processing protocol that enable the establishment of PDTOs and tumor infiltrating lymphocytes (TILs) isolation. By the optimization of functional assays, we compared the T-cells effector functions of matching PBMCs and TILs, demonstrating that PBMCs after co-culture and TILs after initial expansion display similar responses. In addition, the evaluation of cytokine production by fluorospot in combination with an image-based killing assay enable the screening of different immune-checkpoint inhibitors as well as its combination with target inhibitors. Our proof-of-concept functional assays showed the potential and versatility of PDTOs and T-cells co-culture systems for immunotherapy screening. The optimization of scalable functional assays downstream co-culture represents a significant step forward to increase the value of PDTOs as pre-clinical models for immunotherapeutic screens.

## Introduction

Immunotherapy has become a major treatment option in oncology, particularly with immune checkpoint inhibitors targeting the PD-1/PDL-1 pathway^1^. These inhibitors have been approved as first- and second-line treatments, resulting in improved efficacy and longer-lasting responses compared to standard chemotherapy^2–6^. Patients with certain cancers, such as metastatic melanoma and non-small cell lung cancer (NSCLC), have experienced better quality of life and survival rates^2–4, 7, 8^. However, many patients still do not respond or progress within six months of treatment initiation^3–5^. Research is focused on understanding the interaction between tumor and immune cells to identify those who will benefit most from immunotherapy^9–11^.

In NSCLC, PDL-1 expression levels, high PD-1 expressing tumor-infiltrating lymphocytes (TILs), and tumor mutational burden (TMB) are associated with significant responses to anti-PD-1/PDL-1 immunotherapy^12–14^. High TMB tumors may respond better to immune checkpoint therapy when pre-treated with chemotherapy and/or targeted agents^14^. Combining immunotherapy with chemotherapy and/or targeted agents may have synergistic effects, but the safety and efficacy of these combinations are still being explored^11^. Preclinical models are needed to predict treatment sensitivities and guide treatment selection for patients who are unlikely to respond to first-line treatment^15^.

Patient-Derived Tumor Organoids (PDTOs) offer a way to test multiple therapeutic strategies ^16, 17^. They have been successfully used for high-throughput drug screening of chemotherapeutic and targeted agents, but their use in immunotherapy modeling is still limited. Some studies have shown promising results using PDTOs co-cultured with peripheral blood mononuclear cells (PBMCs)^18^ or TILs^19^. However, the optimal method for expanding tumor-reactive T-cells and comparing the killing efficiency of PBMCs and TILs remain to be determined.

Alternative strategies have retained immune components into organoid-like cultures, such as air-liquid interface (ALI) organoids and immune-enhanced organoids, enabling the modeling of patients’ responses to anti-PD-1/anti-PDL-1 treatment^20–22^. However, these models’ scalability and viability for high-throughput drug screening are limited due to variations in cell composition and short-term viability.

To address this, we developed a new tumor processing protocol that enables PDTOs establishment and TILs isolation from the same lung tumor resections. We find that a previously described 14-day co-culture strategy for expanding anti-tumor clones is inadequate for TILs as it leads to an increase in inhibitory receptor expression that impairs their cytotoxic activity. Instead, after their initial expansion they possess comparable or superior cytotoxic activity than matching PBMCs after co-culture with PDTOs. In addition, we also optimized scalable functional assays to evaluate T-cell effector functions by testing various combinations of immune checkpoint inhibitors and target inhibitors. Our results emphasize the immense potential of PDTOs-T-cell co-culture as a highly promising pre-clinical model for immunotherapy screening.

## Results

### A new protocol enables NSCLC-PDTOs establishment and TILs isolation

In order to create PDTOs-T-cell co-cultures, we first sought to establish PDTO and isolate TILs by processing the entire tumor sample with either collagenase I or IV. With collagenase type I treatment, we successfully isolated TILs but could not establish PDTO. With collagenase type IV treatment, we successfully established PDTOs but could not isolate TILs (not shown). Therefore, we used a hybrid protocol depicted in **Supplementary figure 1** where tumor samples are split in half, one half is digested with collagenase IV for the establishment of NSCLC-PDTOs, and the other half is digested with collagenase I for the isolation of lymphocytes. T-cell cultures are monitored until large T-cell clusters are visualized. At that point, a rapid expansion protocol (REP) is started by the addition of irradiated allogenic PBMCs to the TILs cultures (1:200) as feeder layers in addition to IL-2 and α-CD3. After 10 days, TILs samples are frozen down and biobanked. Of note, at the moment of the tumor resection, peripheral blood samples were obtained and processed for PBMCs isolation and cryopreservation. Once the PDTOs reach the growing well status (after several passages and frozen-biobanked vials) molecular characterization and pathological review are performed to evaluate resemblance with the primary tumor (**Supplementary Figure 1**).

Following this protocol, from November 2020 until June 2022, 17 tumor resections were collected from treatment naïve NSCLC-patients, split, and processed for TILs isolations and PDTOs establishment. Patients’ clinical data is depicted in **Supplementary Table 1.** The tumors consisted mostly of invasive adenocarcinomas (LUAD, n=15) and also included 1 squamous cell carcinoma (LUSC) and 1 large cell neuroendocrine carcinoma (LCNEC). PDTOs were successfully established for 9/17 (52.9%) cases, 7 LUAD, 1 LUSC and 1 LCNEC. TILs were isolated and expanded for 7/17 (41.17%) cases. We obtained PDTOs-TILs pairs for 6 patients.

PDTO generation success rate was positively associated with both tumor size and stage but showed no clear association with histological features (**Figure 1A**, **Table 1**). The likelihood of successful TIL expansion moderately increased with tissue size **(Figure 1A)** but, interestingly, it was not associated with the overall level of tumor lymphocytic infiltration (data not shown). Even if tumors showed lymphocytic infiltration, TILs could still fail to expand *in vitro.* Altogether, these results demonstrate feasibility of isolating and culturing matched PDTOs and TILs from the same tumor.

**Figure 1.**
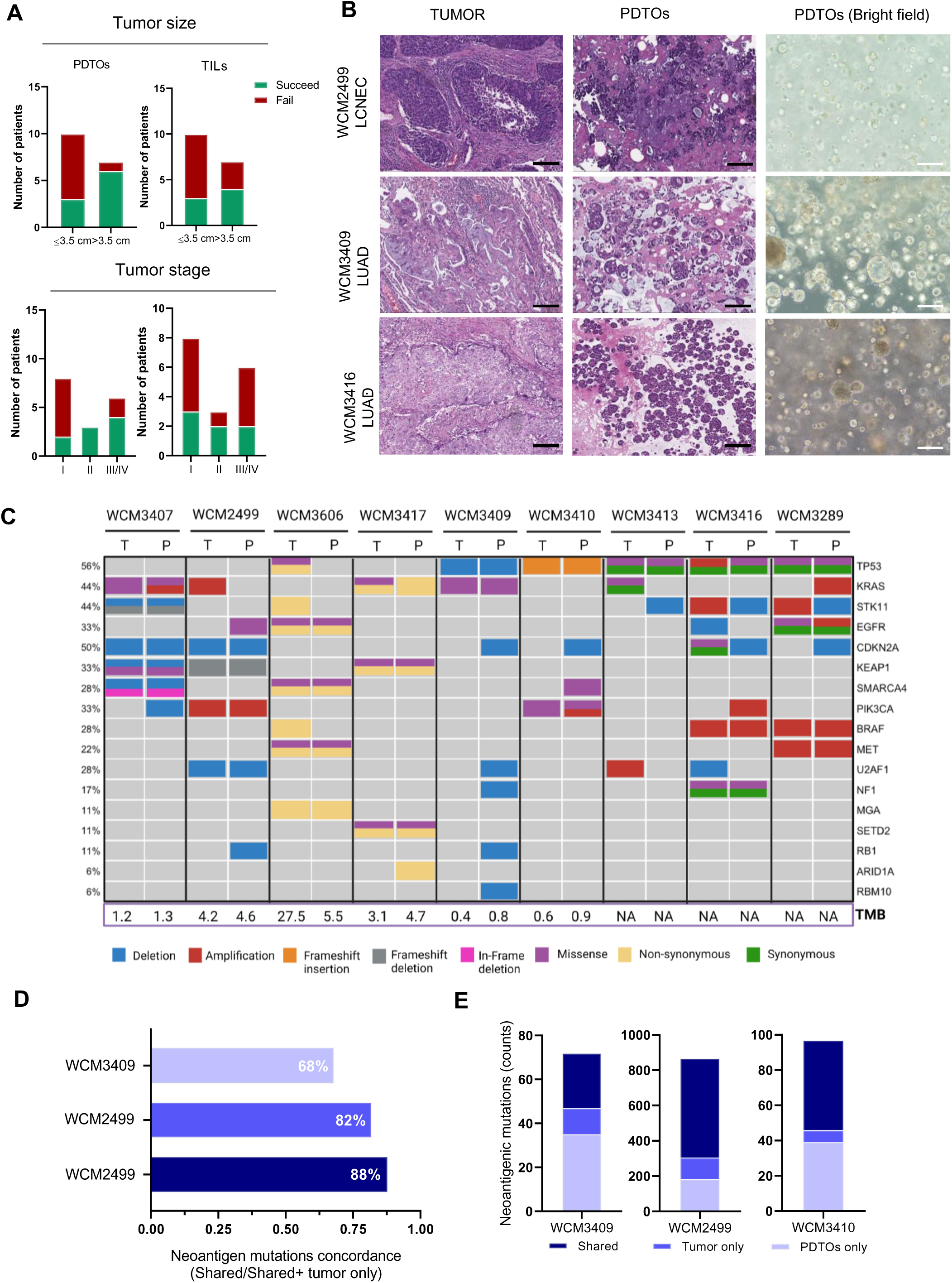
PDTOs establishment and TILs isolation success rate and PDTOs characterization. A. Association of tumor size and disease stage with PDTOs establishment and TILs isolation success rate. Samples were divided according tumor size into low (≤ 3.5 cm) and high (>3.5 cm) and based on disease stage in I, II, III and IV. Number of patients samples that succeed (green) and fail (red) are shown for each group. Fisher’s exact test was used to determine significant associations (p=0.05). B. H&E of tumor and PDTOs pairs for 3 cases of different subtypes of NSCLC: LCNEC (WCM2499), LUAD (WCM3410) and LUSC (WCM3416). Bright filed images for the PDTOs are also displayed. Bars indicate 100μm C. Oncoprint displaying the concordance of molecular alterations in driver genes of PDTOs and tumor pairs. We address the mutational state and copy number alterations of *TP53, KRAS, KEAP1, STK11, EGFR, NF1, BRAF, SETD2, RBM10, MGA, MET, ARID1A, PIK3CA, SMARCA4, RB1, CDKN2A, U2AF1, RIT1, HER2* genes reported to be relevant drivers in NSCLC^31^. Neoantigen-associated mutations were predicted for WCM2499, WCM3409 and WCM3410 based on WES data. D. Neoantigenic mutations concordance for each case calculated as shared mutations/ shared mutations + tumor exclusive mutations. E. Number (counts) of neoantigenic mutations present only in the tumor, only in the PDTOs, or shared.

**Table 1.** Tumor of origin histopathological features and success of PDTOs establishment and TILs expansion. LUAD: Adenocarcinoma; LCNEC: Large neuroendocrine carcinoma; LUSC: Squamous cell carcinoma. * Indicate samples that did not reach the growing well status but enough material was obtained to perform co-culture assays.

| Patient ID | Original histology | WHO 2021 classification | Patterns | PDTOs establishment (YES/NO) | TILs expansion (YES/NO) |
| --- | --- | --- | --- | --- | --- |
| WCM3602 | LUAD | Invasive Non-Mucinous | solid, acinar, lepidic | NO | NO |
| WCM3603 | LUAD | Invasive Non-Mucinous | solid | NO | YES |
| WCM3604 | LUAD | Invasive mucinous | mucinous dominant and micropapillary | NO | NO |
| WCM3605 | LUAD | Invasive Non-Mucinous | acinar, micropapillary, and lepidic | NO | NO |
| WCM3607 | LUAD | Invasive Non-Mucinous | acinar and lepidic | NO | NO |
| WCM3608 | LUAD | Invasive Mixed mucinous and non-mucinous | acinar, solid, micropapillary, lepidic and mucinous | NO | NO |
| WCM3083 | LUAD | Invasive Non-Mucinous | acinar | NO | NO |
| WCM3407 | LUAD | Invasive Non-Mucinous | micropapillary and papillary | YES | NO |
| WCM2499 | LCNEC | LCNEC | LCNEC | YES | YES |
| WCM3606 | LUAD | Invasive Non-Mucinous | micropapillary and acinar pattern | YES* | YES |
| WCM3417 | LUAD | Invasive mucinous | mucinous | YES* | YES |
| WCM3409 | LUAD | Mixed mucinous and non-mucinous invasive | micropapillary, solid, acinar, mucinous and papillary | YES | YES |
| WCM3410 | LUAD | Mixed mucinous and non-mucinous invasive | micropapillary, acinar, mucinous and papillary | YES | YES |
| WCM3413 | LUAD | Invasive Non-Mucinous | micropapillary, acinar and lepidic | YES | NO |
| WCM3066 | LUAD | Invasive Non-Mucinous | solid and micropapillary | NO | NO |
| WCM3416 | LUSC | Invasive SqCC | Invasive SqCC | YES | YES |
| WCM3289 | LUAD | Invasive Non-Mucinous | Solid | YES | NO |

As part of our PDTO quality control pipeline, we performed a pathological review comparing PDTOs and matching tumor histology. Tumor and PDTOs H&Es images for three representative cases of each NSCLC subtype together with bright field images showing PDTOs morphology are shown in **Figure 1B**. The LUSC (WCM3416) and LCNEC (WCM2499) derived PDTOs displayed a similar structure and cell composition to the tumor of origin. Surprisingly, LUAD-derived PDTOs were predominantly acinar and/or mucinous (WCM3409 is shown as illustrative LUAD case) even when the primary tumors patterns were mostly micropapillary suggesting that the culture condition may be promoting a change in morphology.

In addition to histopathology review, we sequenced the tumor and PDTOs pairs and analyzed the concordance of point mutations, indels as well as copy number alterations. We focused our initial analysis on 19 genes previously reported by TCGA as drivers in lung cancer^23^. As reported by TCGA, genomic alterations in *TP53* were the most frequent (56% in our cohort vs 46% TCGA) followed by *KRAS* (44% vs 33%). *KRAS* and *EGFR* were mutually exclusive. In terms of copy number alterations, deletions involving *CDKN2A* were highly frequent (50%) (**Figure 1C)**. Overall, the majority of driver mutations were preserved between PDTOs and tumor of origin (**Figure 1C**).

Whole Exome Sequencing (WES) data was available for 3 of 5 of the cases evaluated in the co-cultures, enabling us to analyze the concordance of predicted neoantigenic mutations between tumor of origin and PDTOs. For the three cases (WCM2499, WCM3409 and WCM3410), most of the neoantigenic mutations found in the tumor were also found in the matched PDTO. The overall concordance was >80% for WCM3410 (88%) and WCM2499 (82%) while it was 68% for WCM3409 (**Figure 1D**). In the latter case, the PDTO presented a larger number of mutations that were not present in the tumor **(Figure 1E).**

### A new optimized approach for assessing the functionality of T-cells in co-culture with PDTOs

We then went on to perform co-cultures for 5 PDTOs-TILs pairs derived in this study. One critical challenge is to assess the functionality of T-cells after co-cultures. Previously, it has been described that co-culture of PBMCs with autologous NSCLC-PDTOs led to an increase in the frequency of tumor reactive IFNγ+ T-cells^18^. In order to assess TILs anti-tumor reactivity (and compare them with autologous PBMCs), we followed the same protocol with minor modifications. Each T-cell sample was divided into two groups, both cultured for 14 days in the presence of IL-2 but only in one group PDTOs were added at day 0 and 7 (co-cultured group). At day 14, T-cells from both groups were harvested, re-plated and co-cultured with PDTOs to perform the functional assays to evaluate T-cell tumor killing capacity and anti-tumor reactivity (cytokine production). Additionally, we added different mAbs targeting immune checkpoints (α-PD-1, α-PDL1, α-PD-1/PDL1 and α-TIM3) at day 0 and 7 of culture, to evaluate their effect during T-cell antigen-specific expansion and once again at day 14 when functional assays were performed (**Figure 2A**).

**Figure 2.**
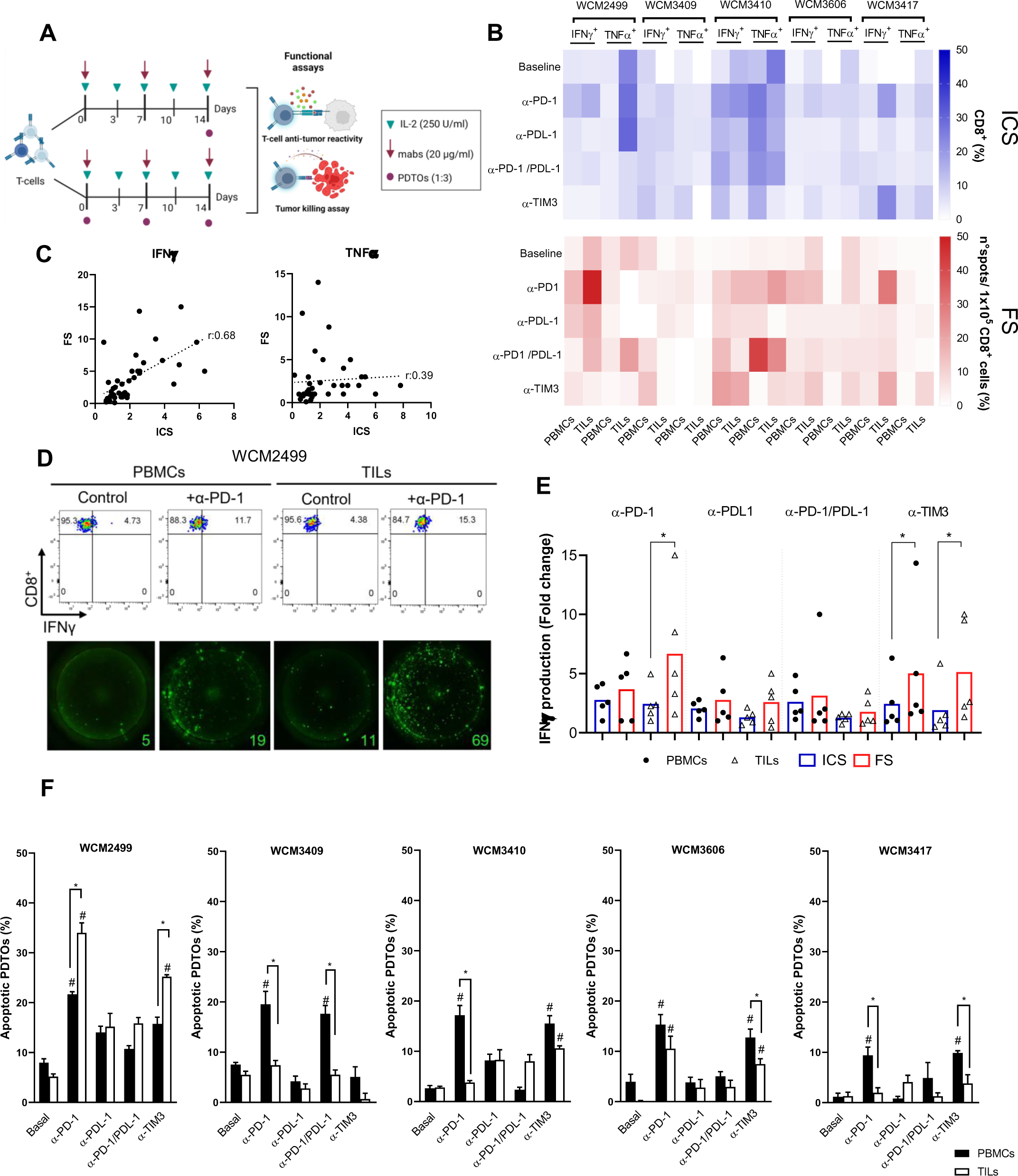
Optimization of functional assays for the evaluation of T-cells effector functions and its modulation by immune-checkpoint inhibitors after co-culture with PDTOs. A. NSCLC-PDTOs and T-cell co-culture protocol. T-cells (TILs or PBMCs) were co-cultured with autologous PDTOs for 14 days. Fresh PDTOs (1:3, Effector: Target) and mAbs (α-PD-1, α –PDL1, α-PD-1/PDL1 and α-TIM3) where added at day 0, 7 and 14. At day 14 T-cells were harvested and functional assays were performed: T-cell antitumor reactivity (IFNγ and TNFα production) and tumor killing assay. B. T-cell antitumor reactivity recorded by intracellular staining (ICS) and fluorospot (FS). Heatmaps displaying the frequencies of T-cells producing IFNγ/TNFα, baseline and upon the addition of different mAbs, are shown. Frequencies are shown as percentage of CD8^+^ IFNγ^+^/TNFα^+^ for ICS (blue heatmap) and as number of spots per 10 x 10^5^ CD8^+^ T-cells for FS (red heatmap). PBMCs and TILs frequencies for each patient are shown (n=5). C. Correlation between IFNγ (left panel) and TNFα (right panel) measures recorded by ICS and FS. Statistical significance was determined by Spearman’s correlation (two tails, CI=95%). Spearman r coefficients are shown for each cytokine. D. Representative dot plots (CD8^+^ vs IFNγ^+^) and FS images (IFNγ spots) for WCM2499 PBMCs and TILs samples, control and with the addition of α-PD-1 (20µg/ml). Percentage of CD8^+^ IFNγ^+^ as well as number of IFNγ spots are depicted. E. Comparison of IFNγ production by T-cells (TILs and PBMCs) in the presence of the different mAbs (α-PD-1, α-PDL1, α-PD-1/PDL1 and α-TIM3) recorded by ICS and FS. Values are shown as fold change relative to the control levels. (*) Statistical significance was calculated by Kruskal-wallis test, Dunn’s multiple comparisons post-test (p=0.05). Black dots: PBMCs, White triangles: TILs, Blue bars: ICS, and Red bars: FS. n=5).F. Image-based tumor killing assay. 3-5 x 10^4^ Far red cell trace stained PDTOs were seeded in 96 well clear bottom plates. T-cells were added (3:1, effector:target) in T-cell culture media containing NucView 488 Caspase-3 substrate (5µM). Cells were co-cultured during 12hs and images were taken every hour. Apoptotic PDTOs (Yellow events)/ field were quantified using Incucyte S3. 3 fields were recorded per well. Apoptotic PDTOs (%) are displayed for PBMCs (black bars) and TILs (white bars) for each patient after 12 hs of co-culture. Results are shown as the average percentage of apoptotic PDTOs (3 fields/condition) ± SEM. Statistical significance was calculated by Kruskal-wallis test, Dunn’s multiple comparisons post-test (p=0.05).

Dijskstra et al reported that the frequency of IFNγ+ T-cells after co-culture increased for most of the cases, yet the overall percentage of these cells was modest (below 5% for most of the patients). We speculated that this could be related to the use of intracellular cytokine staining (ICS) as experimental readout to detect tumor reactive T-cells. Thus, in addition to ICS, we evaluated the production of IFNγ and TNFα using Fluorospot (FS) since this technique is reported to be at least 500 times more sensitive than flow cytometry, enabling the detection of antigen-specific T-cells present in low clonal frequencies^24^. Using both techniques, we observed significant inter-patient heterogeneity in IFNγ and TNFα positivity in T-cells after co-culture with PDTOs, as well as differences in the responses between treatments and between autologous PBMCs and TILs, both at baseline and in response to treatment (**Figure 2B**). A correlation analysis between the T-cell anti-tumor reactivity levels recorded by each technique showed a strong and significant correlation for IFNγ (r=0.68) and a weak (yet significant) for TNF-α (r=0.39) (**Figure 2C**). The difference between the baseline cytokine release and the one recorded in the presence of mAbs was more prominent by FS (**Figure 2D**, bottom row) than for ICS (**Figure 2D**, top row). In addition, fold change values of IFNγ production by PBMCs and TILs after the addition of the different mAbs recorded by FS were higher than those recorded by ICS, especially for TILs with α-PD-1 (2.44 by ICS vs 6.67 by FS) and TILs and PBMCs with α-TIM3 (1.91 vs 5.13; 2.44 vs 5.01) (**Figure 2E)**. Overall, FS yields better resolution (higher fold changes between basal and treated) and higher sensitivity than ICS at evaluating anti-tumor reactivity. This higher sensitivity shown by FS translates into a significantly lower number of cells necessary compared to ICS, which allows to increase the number of experimental conditions that can be assessed for a particular patient sample including multiple drugs combinations.

In addition to the anti-tumor reactivity, we evaluated the cytotoxic activity of the T-cells after co-culture. On day 14, T-cells were challenged with Cell trace Far red-stained PDTOs in T-cell media containing NucView 488 caspase-3 substrate. Pictures were taken every hour using the Incucyte S3 and the apoptotic events were recorded. After 12 hours of culture, the percentage of apoptotic PDTOs in each experimental condition was heterogeneous among the different patients (**Figure 2F).** All patients responded to α-PD-1, increasing the killing (to different extents), followed by α-TIM3, which modulated T-cell responses in 3/5 cases. Of note, with the exception of WCM2499, PBMCs exhibit a higher cytotoxic activity than TILs after co-culture (**Figure 2F**). By optimizing these downstream co-culture functional assays, we were able to systematically evaluate T-cell responses under treatment with different immune checkpoint inhibitors, capturing interpatient heterogeneity, highlighting the value of our experimental platform for immunotherapy screens.

### TILs enable effector functions assessment without long term co-culture with PDTOs requirement

To evaluate how co-culture impacts TILs and PBMCs functionality we compared the cytotoxic activity of the T-cells when co-cultured for 14 days with PDTOs vs those cultured for 14 days alone (just in media supplemented with low IL-2). We found that when co-cultured, PBMCs displayed higher killing levels when α-PD-1 and α-TIM3 were added compared to the basal condition. In contrast, the trend is the opposite for TILs, showing lower basal killing levels after co-culture that are partially reversed with addition of the different mAbs. These results suggest that the 14 days co-culture strategy might not be appropriate when using TILs as the source of T-cells (**Figure 3A**).

**Figure 3.**
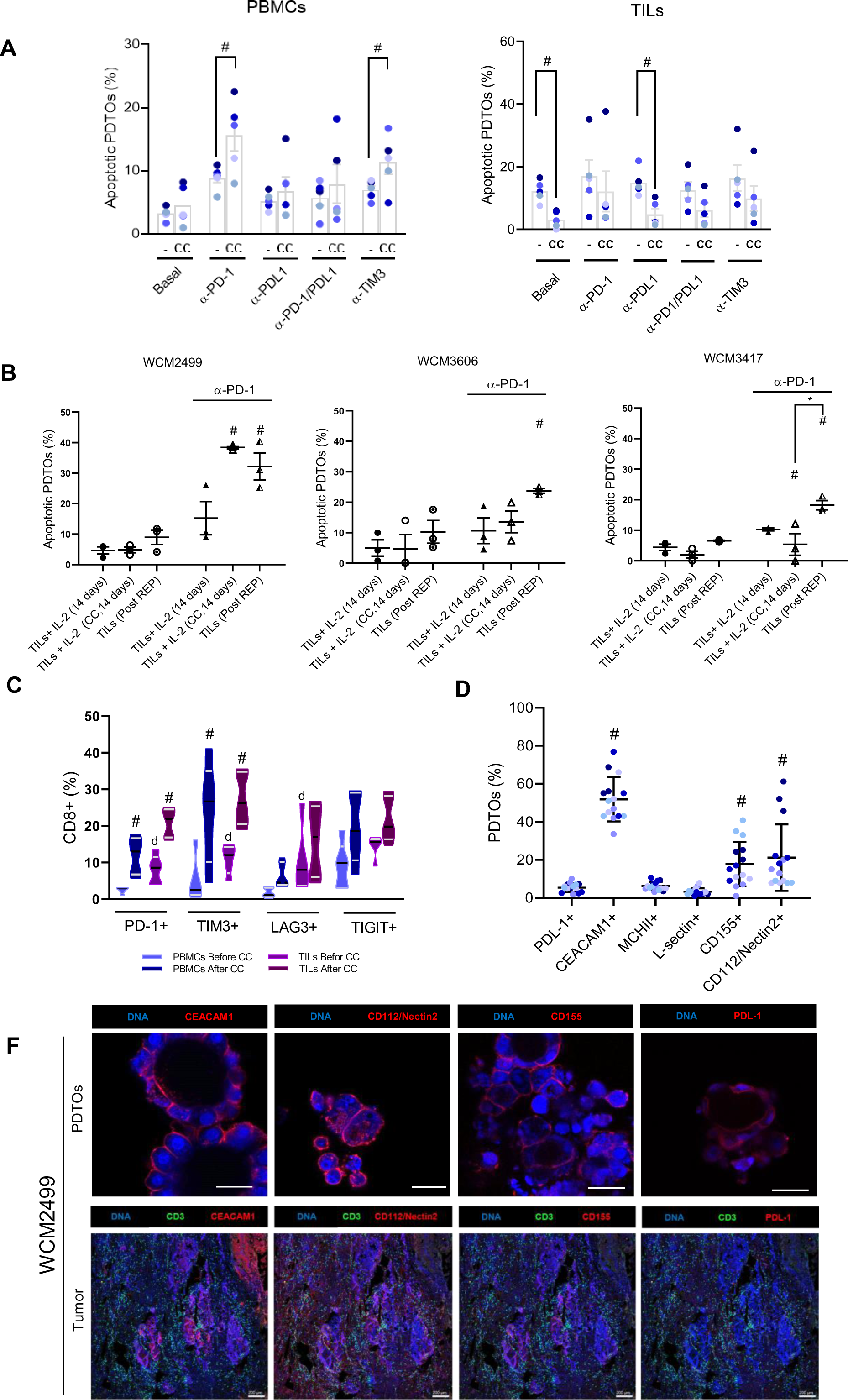
Impact of co-culture on T-cell cytotoxic capacity. A. T-cell mediated tumor killing cultured during 14 days with IL-2 (-) or with IL-2 + PDTOs (co-cultured ; CC) . Apoptotic PDTOs percentage after 12 hours of culture are displayed for each patient in the presence or absence of mAbs. Killing levels recorded for PBMCs (left panel) and TILs (right panel) are shown. # Indicates significantly different. Statistical significance was calculated using Mann-Whitney Test (p=0.05) B. Tumor killing levels recorded for TILs after different culture conditions: Co-cultured for 14 days (CC), cultured for 14 days just with the addition of IL-2 or post REP. Percentage of apoptotic PDTOs are depicted for TILs from WCM1630, WCM1656, and WCM1669 baseline or with the addition of α-PD-1 (20μg/ml). Mean apoptotic PDTOs percentage (3 fields/condition) ± SEM are shown. Statistical significance was calculated by Kruskal-wallis test, Dunn’s multiple comparisons post-test (p=0.05). # Indicate significantly different from the respective condition without α-PD-1. * Indicate significantly different. C. Percentage of CD8+ cells expressing the inhibitory receptors PD-1, TIM3, LAG3 and TIGIT at day 0 and 14. Violin plots displaying the mean (black line) and quartiles (gray lines) for each inhibitory receptor in PBMCs and TILs before and after co-culture. # Indicate different than the condition before co-culture. d indicate significantly different than the PBMCs before co-culture. D. Percentage of PDTOs expressing PDL1, TIM3 ligand CEACAM, TIGIT ligands CD112/Nectin2, CD155, LAG3 ligands MCHII, and LSECtin. PDTOs line were stained 3 times at different passages between passage 6 and 15. # Indicate significantly higher than the PDL1 expression. Differences were determined by Kruskal-wallis test and Dunn’s multiple comparison post-test (p=0.05) E. Representative confocal microscopy images of WCM2499 showing the surface expression of PDL1, CEACAM1, CD112/Nectin2, and CD155. Blue: DNA(DAPI); red: Immune checkpoint ligands (upper row). Imaging cytoff of a tumor resection of WCM2499 showing the overall lymphocytic infiltration (CD3, green), DNA (blue) and expression of PDL1, CEACAM, CD112/Nectin2 and CD155 (red). Scale bars: 200μm (lower row).

In order to confirm the detrimental effect of the 14 days co-culture on TILs cytotoxic activity and based on the premise that TILs constitute a T-cell population already enriched in anti-tumor clones as opposed to their PBMCs counterpart, we compared TIL cytotoxic activity after Rapid Expansion Protocol (REP) with that recorded after 14 days of culture (co-cultured with PDTOs or only with IL-2). We conducted these experiments only with WCM2499, WCM3606 and WCM3417 due to sample availability. Without anti-PD-1 addition, the basal killing capacity of TILs post REP remained as low as the other two culture conditions. However, when α-PD-1 is added during the functional assay, the tumor killing capacity for all the patients increased, both 14-day co-culture (for WCM2499 and WCM3417) and post-REP (all cases). Post-REP TILs achieved highest killing levels in two out of three cases (WCM3606 and WCM3417) (**Figure 3B**). Of note, the cytotoxic activity of the TILs from WCM2499 was not impaired by co-culture and it showed similar levels than the one recorded post REP.

Overall, these results indicate that assessing TILs cytotoxic function after REP improves T cell cytotoxic activity over 14 day cultures, thus confirming that no further culture is required to achieve measurable effector function.

To understand why the co-culture could be affecting T-cell cytotoxic activity, we assessed whether the 14 days co-culture changes the expression levels of inhibitory receptors on T-cells. We evaluated the percentage of CD8+ cells positive for PD-1, TIM3, LAG3 and TIGIT before and after co-culture (**Figure 3C**). After 14 days co-culture, the percentage of CD8+ cells positive for PD-1 and TIM3 increased for both PBMCs and TILs (**Supplementary Figure 2A)**. The initial frequency of CD8+ cells positive for PD-1, TIM3 and LAG3 markers was higher in TILs than PBMCs (**Figure 3C**). By analyzing the expression of inhibitory receptors, we confirmed that the loss of cytotoxic activity on TILs after co-culture is associated with an increase of inhibitory receptors expression. Challenging TILs with PDTOs after initial expansion showed similar results than PBMCs after 14 days of co-culture. These results demonstrate that using TILs offer the possibility to have results in a shorter time frame, increasing the potential value of these models to contribute to the clinical decision making in the future.

### PDTOs recapitulate inhibitory ligands expression profiles of tumors of origin

To better understand which immune checkpoint axes could be impairing T-cells cytotoxic activity in the co-cultures, we evaluated the expression of PDL-1; LAG3 Ligands MHCII and LSECtin; TIM3 ligand CEACAM1; and TIGIT ligands CD155 and Nectin2/CD112, on PDTOs surface by flow cytometry. As expected, the expression profiles were heterogeneous among the different patients, but there were some common features such as the prominent expression of the TIM3 ligand, CEACAM1 (more than 40% of PDTO cells for all PDTOs). TIGIT ligands, CD112/Nectin2 and CD155, were also expressed in a significant percentage of the tumor cells in a less consistent way, the first one ranging between 10 to 55% and the second one between 5 to 40% of PDTO cells. LAG3 ligands displayed a modest (if not absent) expression in most of the PDTO cells (less than 5%) and PDL-1 expression only ranged between 6 to 10% of PDTO cells (**Figure 3D**). Surface expression of PDL-1, CEACAM1, CD112/Nectin2 and CD155 was also evaluated by confocal microscopy. Representative pictures of WCM2499 PDTOs positive for these markers are shown in **Figure 3E upper panel**. Additionally, we assessed the expression of PDL-1, CEACAM1, CD112/Nectin2 and CD155 in the parental tumors to determine if PDTOs recapitulate their matching tumor expression profiles. For that, FFPE tumor fragments were analyzed by Imaging Mass-Cytometry (IMC). In addition to CEACAM1, CD155, CD112/Nectin2 and PDL-1 expression, we assessed CD3 to identify tumor lymphocytic infiltration. LAG3 ligands were not included in the IMC panel due to their low or absent expression on the PDTOs. Tumor staining for WCM2499 is depicted in **Figure 3E lower panel** and images for WCM3409 and WCM3410 are shown in **Supplementary figure 2B.** As observed on the PDTOs, CEACAM1 and CD112/Nectin2 expression was also predominant in their matching tumors, being present in large areas of the tumor sections. CD155 expression was limited on WCM2499 and WCM3409 and completely absent on WCM3410. Of note, PDL-1 expression appears to be minimal in WCM2499 and WCM3410 and absent in WCM3409. Overall, PDTOs recapitulate immune checkpoint ligands representation from their tumors of origin. Considering that CEACAM1 and CD112/Nectin2 were expressed in large regions of the parental tumors as well as represented in high frequency on PDTOs, the combination of TIM3 and TIGIT inhibitors with PD-1 blockade could be a therapeutic strategy for some of these patients.

### Immune-checkpoint inhibitors combinations screening: α-TIM3 and α-TIGIT enhance the effect of α-PD-1 on TILs effector functions

The upregulation of immune checkpoints on T-cells after co-culture as well as the heterogenous expression of its ligands on matching PDTOs make the co-culture systems potentially valuable tools for evaluating combination immunotherapies. As proof of concept, we assessed WCM2499 since the PDTO expressed most of the inhibitory ligands and the TILs displayed significant effector functions (as displayed in Figure 2F). In this assay we combined different concentrations of α-PD-1 with α-TIM3, α-TIGIT or α-LAG3. Two concentrations of the α-PD-1 were assessed (10 and 20µg/ml) with or without the addition of the alternative immune-checkpoints at three different concentrations (5, 10 and 20µg/ml). In agreement with the experiments shown in Figure 2B, we observed that increasing concentrations of α-PD1 boosted anti-tumor reactivity. α-TIM3 increased IFNγ and GrzB activity by itself at 10 and 20μg/ml and it enhanced α-PD-1 effect at both concentrations. α-TIGIT addition only boosted T-cell anti-tumor reactivity at 20μg/ml when α-PD-1 was added (at 20μg/ml). α-LAG3 addition did not have any effect on T-cells reactivity (**Figure 4A).** Representative FS images displaying IFNγ (Upper panel) and GrzB (Lower panel) spots densities for α-TIM3, α-TIGIT and α-LAG3 (20μg/ml), with or without the addition of α-PD-1 (20μg/ml), are shown in **Figure 4B**.

**Figure 4.**
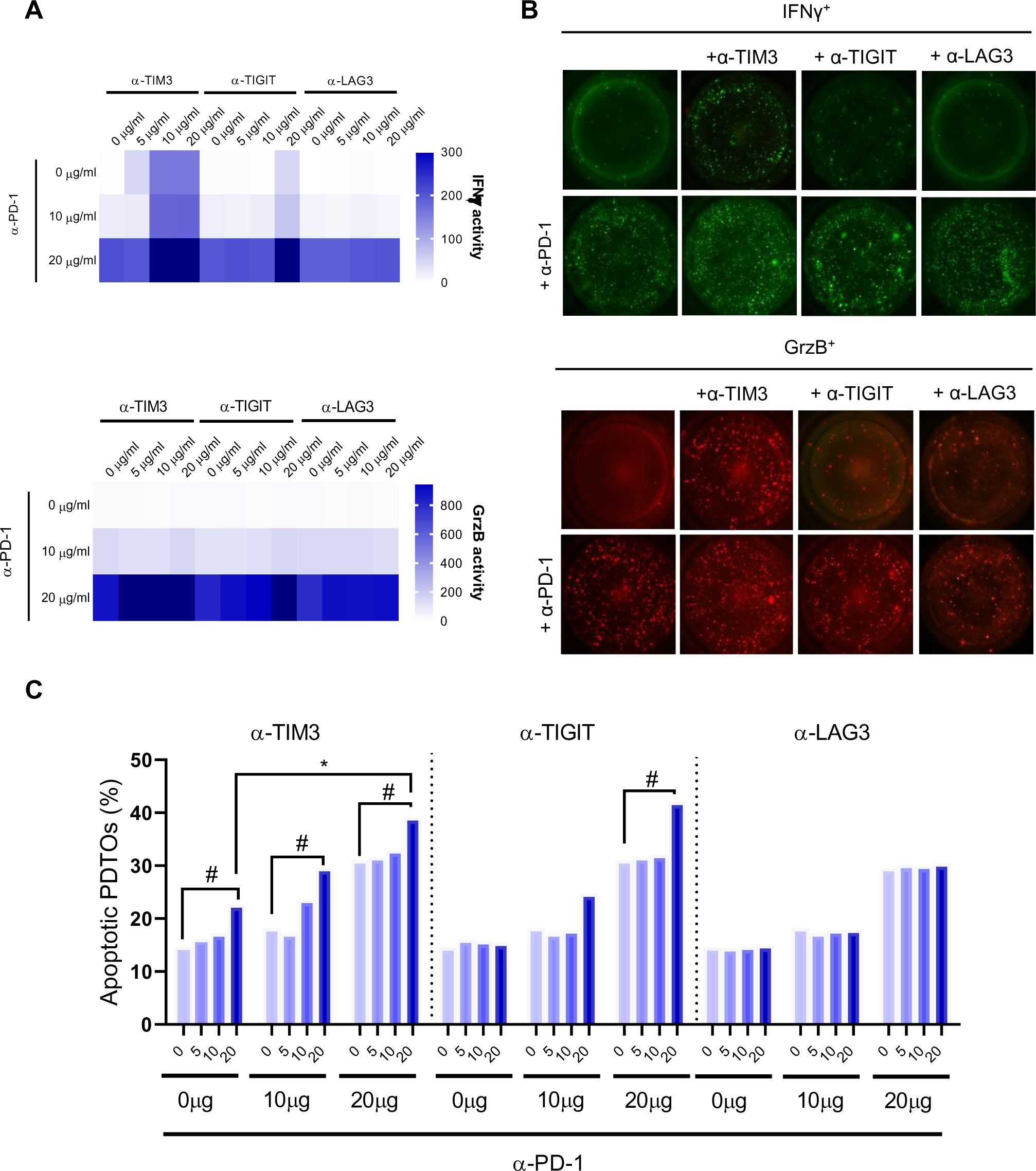
Assessment of immune checkpoint inhibitors combinations. WCM2499 TILs post REP were challenged with PDTOs (1:5) with the addition of α-PD-1 (10-20μg/ml) in combination with α-TIM3 or α-TIGIT or α-LAG3 (5, 10, or 20μg/ml). T-cell antitumor reactivity was measured by FS evaluating IFNγ and GrzB release. A. Heatmaps showing IFNγ and GrzB activity (spots intensity x spots size/1000) values recorded for each combination. B. Representative images displaying IFNγ (green) and GrzB (red) spots density in different experimental conditions. C. T-cell mediated PDTOs killing (percentage of apoptotic PDTOs) recorded for the different immune checkpoint inhibitor combinations. Mean apoptotic PDTOs percentage (3 fields/condition) ± SEM are shown. Statistical significance was calculated by Kruskal-Wallis test, Dunn’s multiple comparisons post-test (p=0.05). # Indicate significantly different from the respective condition with justt α-PD-1 addition. * Indicate significantly different.

The tumor killing assay reflected similar trends as the anti-tumor reactivity assay. The highest concentration of α-TIM3 enhanced T-cell cytotoxic activity in the presence of α-PD-1. With α-TIGIT, T-cell mediated tumor killing significantly increased only in combination with α-PD-1 at 20μg/ml. α-LAG3 addition did not modulate cytotoxic activity (**Figure 4C)**.

These results highlight the potential of the PDTOs/TILs co-culture to address combination of immune checkpoint inhibitors in a patient specific fashion. In this particular case, even when the expression levels of TIM3 and TIGIT (as well as their ligands) were similar, the impact of TIM3 inhibition was more relevant as its addition increased the effects of PD-1 even at suboptimal concentrations (10μg/ml).

### Sequential combination therapy screening: pre- treatment with PI3K inhibitors sensitized *KRAS* (G12A) mutant NSCLC-PDTOs to immunotherapy

Targeted therapies have become a research focal point due to their specificity to tumor cells and minimal adverse effects in comparison to conventional treatments^25^. Nevertheless, a significant number of patients develop resistance that leads to disease progression and death. Current efforts are focusing on new potential combination strategies with synergistic antitumor activity, such as immune checkpoint blockade with targeted agents^11^. However, the identification of effective combinations is difficult due to the large number of possible combinations and the lack of reliable immune-competent models. To establish the utility of our PDTOs/TILs co-culture system for screening combinations of drugs including immune therapies, we performed a drug screening employing two PDTO lines bearing *KRAS* G12A mutations, a low frequency *KRAS* mutation for which there is no clear therapeutic options. Additionally, each of these cases belong to a particular subset of *KRAS* mutants based on the co-occurrence of alterations on *TP53* (WCM3409) or *STK11* (WCM3407). We evaluated if pre-treatment with different targeted inhibitors could sensitize PDTOs to T-cell mediated killing in the presence of immune checkpoint blockers.

As a first step, we evaluated PDTOs sensitivity to 23 different targeted inhibitors against EGFR (Lapatinib, Osimertinib, Dacomitinib, Afatinib, Erlotinib and Gefitinib), PI3K (Idelalisib, Copanlisib, PI-103, Buparlisib, GSK2636771, Parsaclisib and Erganelisib), mTOR (Temsirolimus, Rapamycin and Everolimus), ERK (Ulixertinib), Raf (AZ628 and Dabrafenib), Ras (AMF510) and MEK (Binimetinib, Trametinib and Selumetinib). WCM3409 was sensitive to 7 inhibitors: 4 EGFR inhibitors (Osimertinib, Afatinib, Dacomitinib and Lapatinib), 2 PI3K inhibitors (Buparlisib and Copanlisib) and 1 MEK inhibitor (Trametinib). WCM3407 was sensitive to 9 inhibitors: 5 EGFR inhibitors (Lapatinib, Osimertinib, Erlotinib, Afatinib, Dacomitinib and Gefitinib) and 3 PI3K inhibitors (Copanlisib, PI-103 and Buparlisib). Dose response curves for the hit compounds are shown in **Supplementary Figure 3A.**

To address if these targeted therapies could enhance α-PD-1 and α-TIM3 effect, we tested sublethal drug concentrations (EC10, EC20 and EC25) estimated based on the full dose-response curves (**Supplementary Figure 3B-C)**. To perform these combination assays, we adapted our tumor killing assay. 800-1000 tumor cells were seeded in PDTOs-media Matrigel 25% on 384 wells plates and cultured for 4 days. At day 4 targeted agents were added and incubated for 3 days after which half of the media (15uL) was replaced with T-cell media containing T-cells and the NucView488 caspase-3 substrate. Apoptotic PDTOs were quantified after 24hs of T-cell addition. A schematic experimental timeline is depicted in **Figure 5A**. Fold increase of T-cell killing capacity over the condition without targeted agents is displayed in **Figure 5B-C** for WCM3409 and WCM3407 respectively. In the cases where the fold increase was ≥2 we analyzed differences in the percentages of apoptotic PDTOs to determine if the recorded increases were statistically significant. We found that the addition of EGFR inhibitor, Afatinib, and PI3K inhibitors, Copanlisib and Buparlisib, increased the killing levels in the presence of α-PD-1 and α-PD-1 plus α-TIM3 (5μg/ml) for WCM3409 (**Figure 6B**). The increase in apoptotic PDTOs percentage was particularly pronounced (significantly) for Buparlisib and Copanlisib. In the case of WCM3407, Buparlisib, Copanlisib and PI-103 significantly enhanced the killing percentage when α-PD1 is added alone or in combination with a high concentration of α-TIM3 (**Figure 6C**).

**Figure 5.**
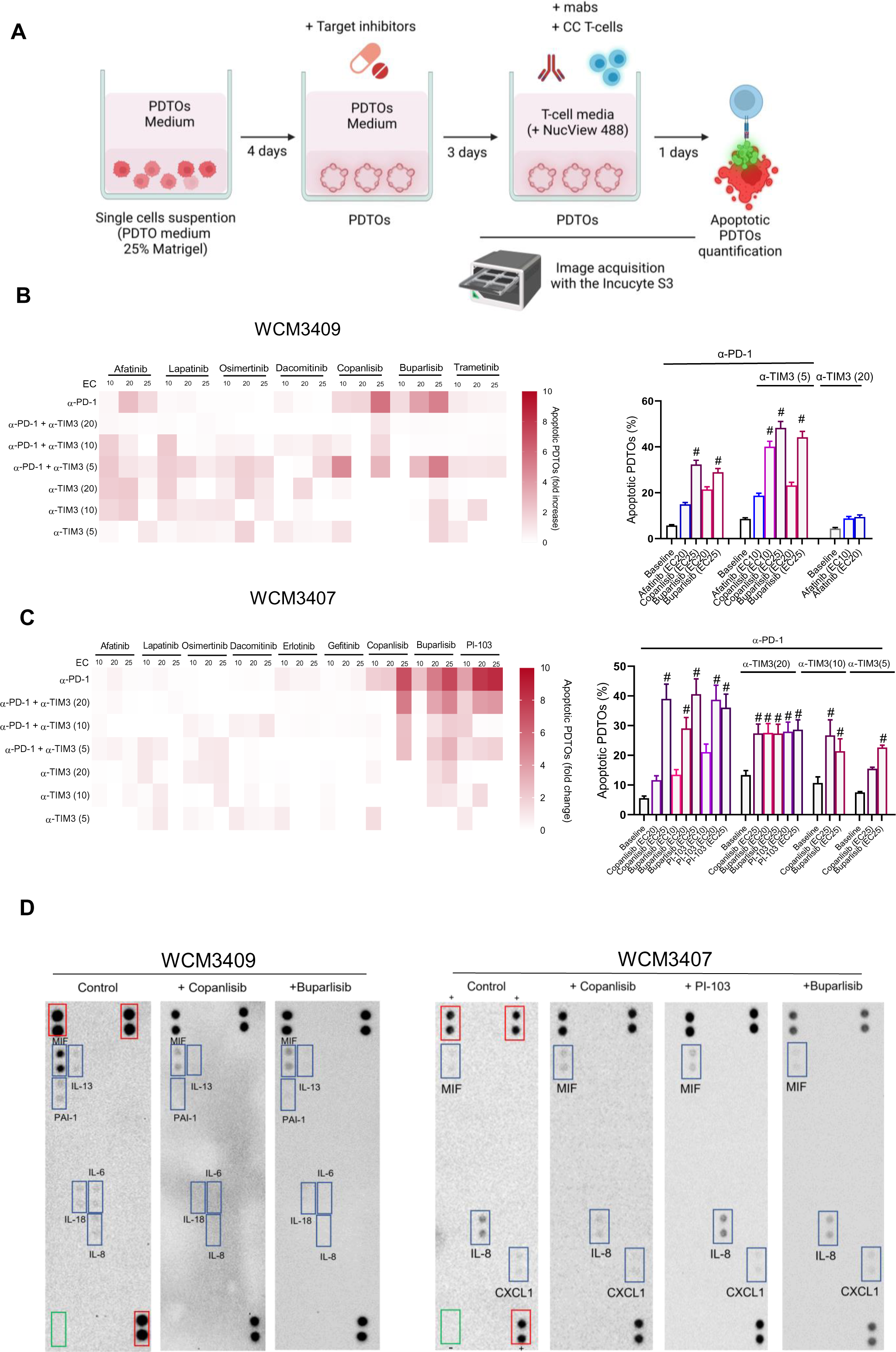
Screening combination therapies for *KRAS* G12A mutants. A. Tumor killing assay adaptation for combination therapy modelling. Far red-stained tumor cell suspensions were plated in Matrigel 25% and incubated for 4 days to enable PDTOs formation. At day 0 target inhibitors were added and PDTOs were incubated for 72hs. At day 7 mAbs and T-cells were added in T-cell media containing NucView488 and imaged O.N using incucyte S3. B-C. Increase in the tumor killing levels mediated by the addition of target agents (EC10, EC20 and EC25). Fold increase was calculated based on the apoptotic PDTOs percentage recorded when only mAbs were added (Left panels). Percentage of apoptotic PDTOs for those conditions where the fold increase was 2 or higher. Mean apoptotic PDTOs percentage (3 fields/condition) ± SEM are shown (Right panels). Statistical significance was calculated by Kruskal-Wallis test, Dunn’s multiple comparisons post-test (p=0.05). # Indicate significantly different from the baseline killing condition. D. Cytokine array of culture supernatants of PDTOs treated for 48hs with the PI3K inhibitors: Copanlisib, Buparlisib and PI-103. Inhibitors were tested at their EC25. Red rectangles: positive controls, green rectangles: negative control, Blue rectangles: cytokines detected.

Since the concentration of the targeted therapies used in these experiments were below EC25, the potentiation on the effect of different mAbs on the PDTOs killing might not be directly related with the exposure of antigens as a consequence of tumor death. In order to find an alternative explanation, we performed a cytokine array (R&D Proteome Profiler Array Cat#ARY005B) on the supernatant of the PDTOs treated with the different PI3K inhibitors to verify if there were variations in the immunoregulatory molecules released by the tumor cells. In both cases we observed that the addition of the PI3K inhibitors reduced the secretion of several cytokines capable of modulate T-cells activity including IL-8 and MIF in both cases, IL-6, IL-13, IL-18 and PAI-1 in WCM3409 and CXCL1 in WCM3407 (**Figure 6D**). These results highlight another advantage of co-culture models in particular of our sequential combination therapy screening, which is the identification of potential microenvironmental signals that could be impairing immune checkpoint inhibitors efficiency leading to the identification of potential new therapeutic strategies.

## Discussion

PDTOs have created a valuable platform for the testing of therapeutic agents and hold great promise for personalized medicine. PDTOs have already been evaluated in the context of high throughput drug screening, modeling clinically applied therapeutic regimens. To assess the predictive potential of these models, patient responses have been compared to the matched PDTOs^26–30^. The results obtained to date indicate that *in vitro* responses of PDTOs are often predictive of patient response to therapy^29, 31^. Further, several groups have developed strategies to maintain or restore immune cells into the PDTO cultures, in order to study the efficacy and specificity of antibody-based therapies targeting aspects of immune function^20, 32^. Nevertheless, there has been no report to date looking to adapt PDTOs cultures to achieve scalable immunotherapeutic modelling.

One of the questions that arises when establishing the experimental design for PDTOs and T-cells co-culture is the source of the latter: PBMCs versus TILs. PBMCs can be easily isolated from peripheral blood samples, and even when the clonality of tumor specific T-cell is low, it can be increased by the co-culture with PDTOs^18, 33^. Previously, it was shown that PBMCs co-cultured with pancreatic PDTOs showed a selection of tumor specific T-cell receptors shared with those present in the TILs populations^33^. In line with that observation, we demonstrated here a high preservation of predicted neoantigenic mutations between tumor and matching PDTOs. The presentation of peptides derived from these mutated proteins in the context of MHC class I molecules in the surface of the PDTOs could lead to the expansion of neoantigen-specific CD8+ T-cells. In our cohort, the cytotoxic activity recorded for WCM2499 TILs were significantly higher than the other cases in concordance with the fact that this tumor/PDTOs pair has ten times more neoantigenic mutations than the other cases (shown in figure 1E). This observation highlights the potential of these co-culture strategies for the identification and expansion of neoantigen-specific clones that could be employed in cell-based immunotherapy. TILs have been used for cell therapy approaches for several types of solid tumors based on the reasoning that they are enriched in tumor reactive T-cells^34, 35^. Their value as a predictive biomarker of immune checkpoint blockade response is being evaluated in several types of solid tumors including NSCLC^12, 36^. Here we evaluated the antitumor functions of TILs and PBMCs after co-culture with autologous NSCLC-PDTOs. We followed the co-culture protocol described by Dijkstra et. al, incorporating different checkpoint inhibitors to evaluate if they can modulate the expansion of tumor reactive clones in both sources of T-cells. In an effort to enable the scalability of the functional assays downstream of co-cultures, we compared INFγ and TNFα production evaluated by ICS with FS. We confirmed that FS achieves higher resolution and sensitivity, using 10 times lower cellular input than ICS.

We also uncovered that cytotoxic activity of TILs was impaired by the co-culture, even when IFNγ/TNFα release levels for TILs and PBMCs were similar. This decrease in the cytotoxic activity of TILs was associated with a higher expression level of exhaustion molecules (TIM3, PDL1, LAG3 and TIGIT). During the process of TIL isolation and expansion, T-cells undergo several rounds of amplification in response to polyclonal stimuli such as IL-2 and α-CD-3. Furthermore, as part of the traditional procedures of TIL manufacturing, they are subjected to a selection phase where T-cells are assessed for specific tumor recognition, in our case, the 14 days co-culture with autologous PDTOs. This whole process makes TILs prone to exhaustion limiting their functionality^37, 38^. Alternative approaches are being evaluated to improve TILs obtention protocols such as the replacement of IL-2 for IL-15 and IL-21 and avoiding selection steps in vitro by directly employing post -REP TILs, called young-TILs^35, 39^. Following this approach, we found that TILs post REP display a higher cytotoxic activity than their after-co-cultured counterparts. Since the cytotoxic levels recorded by TILs after REP and PBMCs after co-culture were similar, the decision of which source to use for the assays will depend on the tumor type and the type of sample, biopsy or resection. We encourage the use of TILs, if possible, because the translational implication of the screening would be more valuable and the timeframe required to obtain data would be significantly shorter. Nevertheless, understanding the low success rate of their obtention, expanding tumor reactive T-cells from PBMCs remains a reasonable alternative.

Currently, PD-1 blockade is approved for use in first- and second-line treatment in advanced non-squamous NSCLC^40^. Unfortunately, many NSCLC patients do not respond or do so only briefly and then relapse. The mechanisms behind this are being extensively investigated. One major known mechanism is the expression of other inhibitory receptors on TILs after the initial upregulation of α-PD-1^41–44^. As an increasing number of checkpoint molecules are discovered on exhausted T-cells, it is essential to understand which of them has dominant effect to design effective combination strategies that can become increasingly complex with significant toxicity profiles. To achieve this goal, robust human based pre-clinical models are needed. The expression of different exhaustion molecules on TILs post REP and on PBMCs after co-culture offers an opportunity for the evaluation of immune checkpoint inhibitor combinations. Among the exhaustion markers that we evaluated in our cohort of patients, TIM3 and TIGIT were the receptors with highest expression levels. Previously, the combinations of α-PD-1 with α-TIM3 or α-TIGIT were assessed in a restricted human TILs adoptive transfer model employing an engineered lung cancer cell line (A549) expressing NY-ESO and HLA-A2 (A549-A2-ESO) and the Ly95 T-cell expressing NY-ESO-1 TCR^45^. In that model, the authors reported that the combination of α-PD-1 either with α-TIM3 or α-TIGIT reinvigorates Ly95 T-cells reactivity and tumor control^45^. Using our co-culture platform, we were able to achieve similar results employing WCM2499 TILs after REP and autologous PDTOs. The combination of α-PD-1 either with α-TIM3 and α-TIGIT increased PDTOs killing and IFNγ and GrzB release. Of note, here we only explore well known immune-checkpoint receptors, but combining these models with next generation sequencing strategies and molecular editing tools, would enable the identification of potential new molecules that could be targeted to potentiate current therapies or develop new ones.

The identification of subsets of patients with oncogenic drivers has transformed the treatment of NSCLC, particularly for those patients whose tumors harbor mutations in EGFR or fusions involving the ALK, RET, and ROS1 kinases^46, 47^. However, these genomic alterations occur in a relatively small percentage of NSCLC patients, mainly LUAD, and when actionable, the efficacy of the available targeted drugs is limited due to the development of acquired resistance through different molecular mechanisms^48^. Therapeutic approaches combining target inhibitors and immunotherapies are being evaluated for these subsets of patients constituting a promising therapeutic field^11^. Nevertheless, the development of therapeutic strategies for *KRAS* mutants, the most common oncogenic driver in NSCLC, has not been successful so far^49^. The development of effective therapies for *KRAS* -mutant NSCLC is challenging due to the heterogeneity in their biology and therapeutic responsiveness^49^.

Besides the alteration in *KRAS*, the co-occurrence of other genomic alterations in *TP53, STK11* and *CDKN2A/B* define particular clusters of patients (named KP, KL and KC respectively)^49^. One of the many molecular differences between the three clusters is the signature of immune-related genes. GSEA and Ingenuity Pathway Analysis highlighted gene sets associated with activation of antitumor immunity and immune tolerance/ escape as prominent modules of the KP cluster. In contrast, KL tumors demonstrated a comparative lack of immune system engagement, whereas KC tumors demonstrated a mixed picture^49^.

In a proof-of-concept experiment, we used WCM3409 and WCM3407, both LUAD carrying a low frequency *KRAS* mutation (G12A) and belonging to the KP and KL clusters, respectively. In an initial screening phase, we evaluated their sensitivity to a set of 23 inhibitors targeting EGFR, PI3K, mTOR, ERK, Raf, Ras and MEK. Both lines were mostly sensitive to EGFR and PI3K inhibition. In this experiment, we tested a sequential treatment strategy, where the PDTOs were first treated with sublethal concentrations of EGFR and PI3K inhibitors for 3 days and then T-cells were added in the presence of mAbs for an additional day. Our aim was to determine if pre-treatment with the inhibitors sensitized PDTOs to T-cell killing. PI3K inhibitors enhanced killing mediated by α-PD-1 and α-TIM3 in both cases.

It has previously been reported that PI3K inhibition increases immunotherapy effectivity, either acting directly on the tumor cells making them more sensitive to immune recognition or enhancing T-cell effector functions^50–52^. In addition, *KRAS* mutations have been associated with tumor promoting inflammation and with the secretion of immune-suppressive cytokines^53, 54^. Considering this, we evaluated in our sequential co-culture system if the enhancement on immune-checkpoint inhibitors effect on T-cell mediated tumor killing when PDTOs were pre-treated with PI3K inhibitors was related with the modulation of the cytokines profile secreted by the tumor cells. Analyzing the supernatant of the PDTOs treated with the PI3K inhibitors for 48hs we observed that both patients shared a common alteration, the reduction of the amount of IL-8 released by tumor cells. IL-8 not only promotes NSCLC cells growth and survival but also interferes with T-cell function by inducing the upregulation of PDL-1 on tumor cells and inducing the apoptosis of subsets of effector CD8+ T-cells^55, 56^. In addition to IL-8, IL-6 and MIF are the only cytokines detected that could be contributing to impairing T-cell functions in our system since they are capable of directly affecting T-cells effector function^57–59^. IL-13, CXCL1 and PAI-1 impair T-cell activation and immunotherapy effectivity indirectly by the interaction with myeloid cells subsets which are not present in our *in vitro* system, thus, their involvement might not be relevant to explain our observations^60–62^.

In this manuscript we described a processing protocol to obtain TILs and PDTOs from the same tumor resections as well as the optimization of scalable downstream co-culture functional assays that enable the assessment of multiple experimental conditions. We demonstrated that our co-culture strategy is suitable for the testing of combination therapies, not only of different immune-checkpoint inhibitors but also their combination with targeted inhibitors.

PDTOs as pre-clinical models are still in an early stage for those therapies where the presence of immune or stromal cells is essential. Validating PDTO responses with patient’s responses is imperative to confirm if these co-culture platforms are significantly useful as preclinical models for immunotherapy. There is still a long road ahead and room for improvement, but there is no doubt of the positive impact that these models will have in precision cancer immunology in the upcoming years.

## Materials and Methods

### Culture media formulations and reagents

Culture media formulations and the list of reagents employed are shown in Supplementary file 1.

### Patients samples

Tissue and whole-blood specimens were obtained from 17 patients with lung cancer diagnosis that underwent surgical procedures between November 2020 and June 2022. Specimens were obtained in accordance with New York Presbyterian hospital-Weill Cornell Medical College (NYPH-WCM) guidelines and under an Institutional Review Board approved protocol (IRB #1008011221). We only included tumors that were larger than 2cm. Once resected, tumor samples were placed on transport media and kept on ice until processing (within 5 hours after resection). A small fraction of the tumor resection was placed in formalin for subsequent histopathological review. PBMC samples were retrieved from the NYPH-WCM thoracic surgery biobank at the moment of the co-culture experiments.

### NSCLC-PDTOs establishment and TILs isolation

When received, tumor resections were split in two pieces. One half was processed for PDTO establishment and the other for TILs isolation.

PDTOs were developed as previously described by Pauli et al. with modifications^16^. Fresh tissue samples were washed three times with transport media and placed in a sterile 3-cm petri dish for mechanical dissection into smaller pieces (2mm diameter) prior to enzymatic digestion. Media containing loose cells or clumps of cells after mechanical dissection were separated from tissue pieces as a “pre-digest” fraction and used later for culture without enzymatic digestion. Enzymatic digestion was done with collagenase IV media in a volume of at least 20 times the tissue volume and incubated on a shaker at 200 rpm at 37°C until the digestion solution turned cloudy, typically 30-45 min. The suspension and the pre-digest fraction were both centrifuged at 300g for 3 minutes and the cell pellet was washed once with washing media. The cells in each fraction were resuspended separately in a small volume of PDTO culture media. Up to ten 100 μl drops of Matrigel/cell suspension were distributed into a 6-well cell suspension culture plate (Cellstar cat #657185). The drops were allowed to polymerize for 30 min inside the incubator at 37°C and 5% CO2 and afterwards, 3 ml PDTO culture media were added per well. Fresh culture media was replaced every 3 to 4 days. PDTOs at approximately 300 to 500 μm were passaged using TrypLE Express for 10-12 minutes in the water bath at 37°C. Single cells and small cell clusters were replated according to the procedure described above. Monthly mycoplasma screening was performed using the abm® Mycoplasma PCR Detection Kit (Cat# G238). PDTOs were cryopreserved in Recovery Cell Culture Freezing Medium (Cat# 12648010) in liquid nitrogen.

For TILs isolation, samples were mechanically dissected, resuspended in Collagenase I media and incubated for 30 min at 37°C in a humidified incubator. After that, digested tissue samples were chopped again and incubated in Collagenase I media for another 30 minutes. After the incubation, digested tissue was chopped again, and cellular suspensions were filtered through 40µm cell strainers and centrifuged at 300g for 3 minutes. Cells were resuspended and incubated for 3 minutes in ACK lysis buffer (Quality biological cat #119-156-721) to eliminate contaminant erythrocytes, centrifugated again and resuspended at a concentration of 5x10^5^ cells/ml in T-cell culture media supplemented with 3000u/ml IL-2 and 0.5 μg/ml anti-CD3 (OKT3 clone) to favor initial T-cell proliferation. Half of the media was replaced every 3 days.

### T-cell rapid expansion protocol

Once T-cell clusters were observed indicating that the cultures were stabilized, cells were harvested and a rapid expansion protocol (REP) was performed to increase the number of available T-cells. REP was performed as previously described by Jin et al.^63^. Briefly, stabilized TILs were cultured with irradiated allogeneic PBMCs (40 or 50Gy) at a ratio of 1:200 in T-cell culture media supplemented with 3000u/ml IL-2 and 1 μg/ml anti-CD3 (OKT3 clone), refreshing half of the media every 3 days. After 14 days of culture, cells were harvested and cryopreserved on T-cell freezing media.

### NSCLC-PDTOs histopathological and genomic characterization

PDTOs histopathology was verified by comparing sections from formalin-fixed and paraffin-embedded (FFPE) passage 5 PDTOs blocks to parent tumor sections using our developed cytology and histology platforms^64, 65^. Briefly, PDTOs were released from Matrigel droplets using cell recovery solution, suspended in a fibrinogen/thrombin gel pellet, fixed with 4% paraformaldehyde in PBS, and embedded in paraffin to create FFPE blocks. Hematoxylin and eosin (H&E) stained sections of the FFPE blocks were verified as tumor cells and compared to H&E-stained sections from the corresponding tumors to verify matching cellular morphology by a WCM pathologist. Whole exome sequencing (WES) or Oncomine/ TruSight Oncology (TSO) 500 targeted sequencing was performed on PDTOs pellets from passage 5 and matching tumors to confirm identity and mutational profile concordance. Single nucleotide variants found in tumor and PDTOs samples via WES, Oncomine or TSO 500 were compared in order to verify concordance of driving mutations in matching samples. We addressed the mutational state and copy number alterations of *TP53, KRAS, KEAP1, STK11, EGFR, NF1, BRAF, SETD2, RBM10, MGA, MET, ARID1A, PIK3CA, SMARCA4, RB1, CDKN2A, U2AF1, RIT1, HER2* genes reported to be relevant drivers in NSCLC^23^.

### Prediction of neoantigenic mutations

To identify potential neoantigens from non-synonymous and truncating mutations pVACtools were used ^66^. Seven epitope prediction methods (MHCflurry, MHCnuggetsI, NetMHC, NetMHCpan, PickPocket, SMM, and SMMPMBEC) were applied to identify neoantigens restricted to patient specific MHC Class I alleles (HLA-A, -B, -C). For each prediction, method with best binding affinity (i.e., lowest IC50) was used to filter and prioritize neoantigens as follows: 1) Neoantigens with IC50 <= 1,000 uM, 2) Fold change between mutant vs corresponding wild-type epitope >=2, and 3) when available, gene expression value (FPKM) of > 1.

### T-cell and NSCLC-PDTOs co-culture

As previously described by Djskstra, et al; tumor reactive T-cells were expanded by co-culturing them during 14 days with autologous PDTOs in T-cell culture media supplemented with 300UI/ml of IL-2. T-cells were rechallenged at day 7 of co-culture and half of the medium was refreshed every 3 days. At day 14, T-cells were rechallenge with PDTOs and functional assays were performed. Different monoclonal antibodies (mAbs) provided by Eli Lily: anti-PD-1 (LSN3415244), anti-PDL1 (LY3300054), anti-TIM3 (LY3321367) and anti PD-1/PDL1 (LY3434172) were added at day 0, 7 and 14 of co-culture to address their impact on tumor specific-T-cell expansion and effector function. Matching isotype huIgG were used as controls.

### Quantification of IFNγ, TNFα and Granzyme B production by effector CD8+ T-cells

#### a. Intracellular cytokine staining (ICS)

After 14 days of co-culture with PDTOs, T-cells were collected and seeded in round bottom 96-well plates. PDTOs were added in a 1:5 (PDTOs: effector cell) ratio. After 1 hr of incubation, GolgiStop (BD) was added (to allow intracellular cytokines accumulation) for 4hs. After this incubation, cells were harvested and washed once with PBS. Anti-CD3, CD4 and CD8 surface stainings were performed by incubating cells with the respective antibody cocktail for 20 minutes. Cells were washed, stained with the fixable viability dye during 20 min at room temperature. Then, cells were washed, fixed and permeabilized with BD fix and perm kit following manufacturer instructions. After permeabilization, cells were incubated with anti-IFNγ, anti-TNFα and anti-Granzyme B for 20 minutes. Finally, cells were washed with PBS and seeded in FACs tubes for acquisition employing BD symphony A5 cytometer.

#### b. Fluorospot Assay (FS)

T-cells were collected after 14 days of co-culture and samples were enriched on CD8+ T-cells using the mojosort CD8+ negative selection Kit following the manufacturer instructions. 100.000T-cells/well were seeded in pre-coated FS plates (mabtech FluoroSpot flex, IFNγ-TNFα-GrZB kit) in 200ul of T-cell culture media and incubated over night with PDTOs in a 1:5 ratio. As positive controls, T-cells were stimulated with Phorbol 12-myristate 13-acetate (25ng/ml) and Ionomycin (1µg/ml). To record basal detection levels, T-cells were seeded in T-cell media alone. FS plates were developed following manufacturer instructions. Plates were sent to Zellnet Consulting for reading services (Mabtech IRIS). Number of spots and activity (spots intensity x spots size/1000) were reported for each fluorescence channel.

### PDTOs killing assay

T-cells recovered after 2 weeks of co-culture were rechallenge with PDTOs to assess their cytotoxic potential. 3 days before the tumor killing assay, 3-5 x 10^4^ single tumor cells were stained with Cell trace far red and seeded in 100ul Matrigel 66% droplets. After 72hs far red stained PDTOs were harvested using cell recovery solution and seeded in 96 well plates in 150 µl of T-cell media containing 5uM NucView488 caspase-3 substrate. T-cells were added in a 3:1 ratio (effector:target) in 50 µl of media. Plates were imaged every 1 hr using Incucyte S3 (Sartorius) for 12 hr recording 3 fields/well. A minimal threshold area of 50μm^2^ was established for red events (PDTOs) to be quantified. Apoptotic PDTOs were identified as double positive cells (green + red signal). Percentage of apoptotic PDTOs was calculated based on the total number of red events/field. Representative cell masking setting for the identification of apoptotic PDTOs and assay images are shown in **Supplementary figure 4**

### PDTOs drug sensitivity assays

PDTOs were digested into a single cell suspension and cells were plated in a 384 well plate (Thermo Scientific Nunc cat#142761) at a density of 1000 cells per well in 8ul droplets (1:2 media:matrigel). Plates were centrifuged briefly to ensure the cells were at the bottom of the well and 15μl of media were added. Cells were incubated for 72h to allow the cells to form PDTOs and afterwards the drugs were added and incubated for additional 96h. The readout was performed using CellTiterGlo®3D reagent according to the manufacturer’s protocol. Luminescence was measured by the Biotek Synergy H4 plate reader.

### Tumor slides imaging mass-cytometry

Banked human lung FFPE tissue samples were baked for 2 hours on a slide warmer at 60 ^0^C. The slides were then dewaxed in CitriSolv (Decon Labs, cat#1601) twice, each for 10 minutes, followed by hydration in descending series of ethanol (100%, 95%, 80% and 75%) for 5 minutes each. The slides were washed in Milli Q water and processed for antigen retrieval using Heat Induce Epitope Retrival method at 95^0^C for 30 minutes^67^. The slides were then cooled to room temperature and washed twice in TBS. After blocking the slides for an hour in SuperBlock^TM^ blocking buffer (Thermo Fisher cat#37515) the slides were incubated in antibody cocktail solution (Antibody Panel described in Supplementary file 1) overnight in 4^0^C. Next day, the slides were washed in 0.2% Triton-X 100 solution twice, TBS twice and incubated in Iridium DNA solution for 30 minutes in RT. The slides were air dried before setting up on the instrument. Imaging Mass cytometer was tuned before setting up the slides and the region of interest was drawn using H&E reference annotation. Images were processed using MCD Viewer software.

