## Supplementaty Figures for "Novel co-culture strategies of tumor organoids with autologous T-cells reveal clinically relevant combinations of immune-checkpoint and targeted therapies"

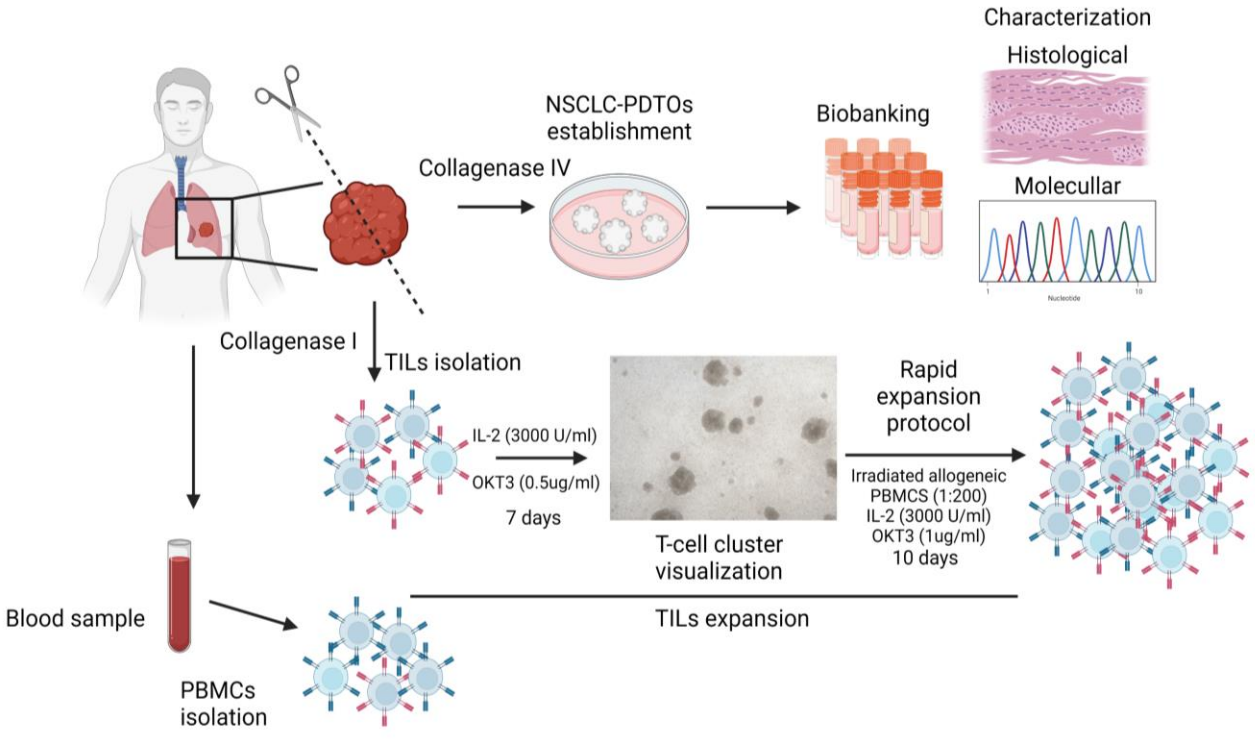


**Supplementary figure 1. Tumor processing protocol for the establishment of PDTOs and TILs isolation.** NSCLC tumor resections that met the inclusion criteria (tumor size ≥ 2cm) were processed for PDTOs establishment and TILs isolation following our processing protocol depicted in **A**. For PDTOs generation tumors were digested with collagenase IV, single cells suspensions seeded in matrigel domes, expanded and biobanked. After PDTOs establishment histopathological review and molecular characterization were performed to evaluate concordance with the tumor of origin. For TILs isolation tumor fragments were digested with collagenase I. Single cells suspensions were plated in T-cell media supplemented with 3000UI/ml IL-2 and 0.5µg/ml α-CD3. When T-cell clusters were observed in the cultures (~7days) clusters were disaggregated, cells harvested, and Rapid Expansion Protocol (REP) was performed to increase the number of cells available for testing. For each patient, peripheral blood samples were collected at the moment of tumor resection and PBMCs were isolated.

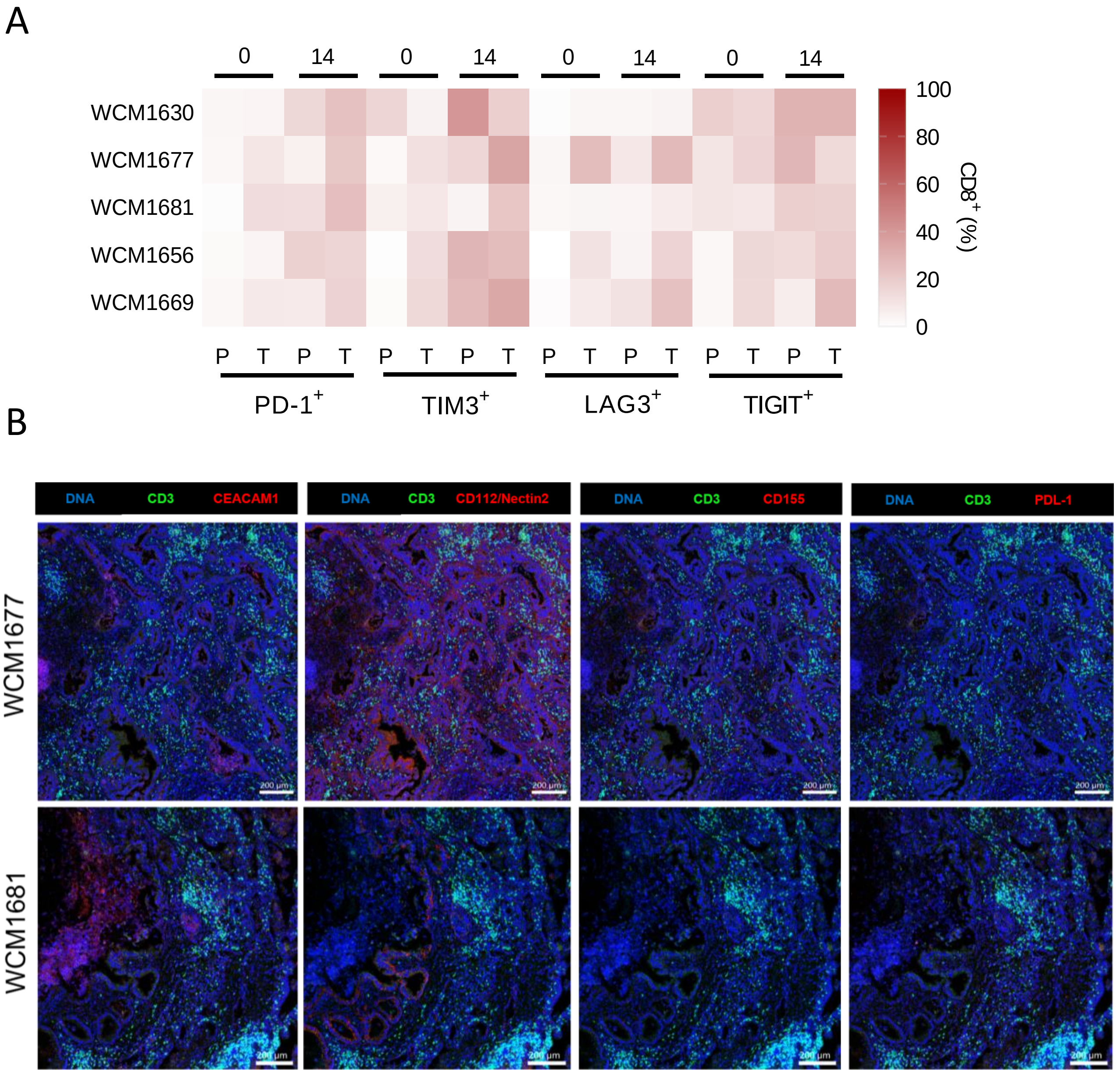


**Supplementary Figure 2. Assessment of immune-checkpoint moleculles on T-cells and tumors.**
**A.** Percentage of CD8+ cells expressing the inhibitory receptors PD-1, TIM3, LAG3 and TIGIT at day 0 and 14. PBMCs (P) and TILs (T) samples are shown for each timepoint and patient.
**B.** Expression of inhibitory ligands and lymphocytic infiltration in tumor sections. Tumor tissue FFPE slides were stained to assess CEACAM1 (TIM-3 ligand), CD112/Nectin2 and CD155 (TIGIT ligands) and PDL-1 expression (red). In addition, α-CD3 was added to analyze the overall lymhocytic infiltration (green). DNA was stained to display tissue structure (blue). Scale bar indicate 200µm. Staining for WCM1677 (upper row) and WCM1681 (lower row) are shown.

A

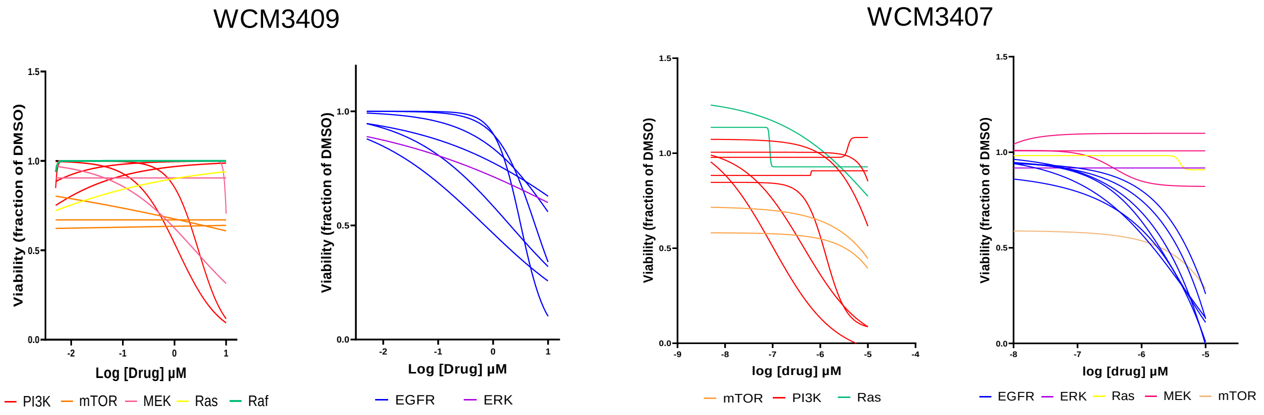


B

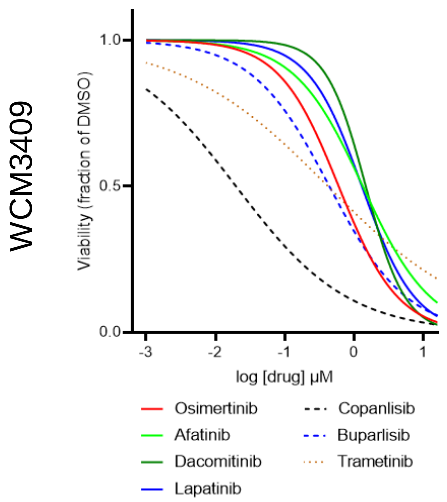


| | EC10 | EC20 | EC25 | EC50 |
|---|---|---|---|---|
| **Afatinib** | 0.268 | 0.5985 | 0.800 | 2.394 |
| **Lapatinib** | 0.431 | 0.47 | 0.9 | 1.877 |
| **Osimertinib** | 0.064 | 0.18 | 0.216 | 0.726 |
| **Dacomitinib** | 0.03 | 0.5 | 1.1 | 2.0 |
| **Copanlisib** | 0.005 | 0.016 | 0.021 | 0.064 |
| **Buparlisib** | 0.008 | 0.017 | 0.398 | 0.7655 |
| **Trametinib** | 0.015 | 0.032 | 0.08 | 0.136 |

C

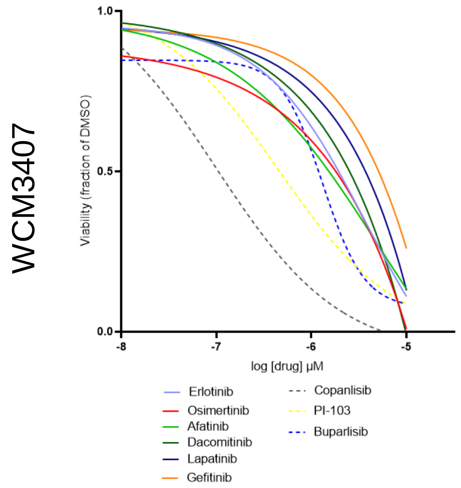


| | EC10 | EC20 | EC25 | EC50 |
|---|---|---|---|---|
| **Afatinib** | $5.85 \times 10^{-7}$ | $1.32 \times 10^{-6}$ | $1.32 \times 10^{-6}$ | $5.3 \times 10^{-6}$ |
| **Lapatinib** | 0.044 | 0.09 | 0.132 | 0.398 |
| **Osimertinib** | 4.13 | 9.3 | 12.4 | 37.20 |
| **Dacomitinib** | 0.07 | 0.162 | 0.216 | 0.6485 |
| **Erlotinib** | $5.85 \times 10^{-7}$ | $1.32 \times 10^{-6}$ | $1.32 \times 10^{-6}$ | $5.3 \times 10^{-6}$ |
| **Gefitinib** | $1.163 \times 10^{-5}$ | $2.62 \times 10^{-5}$ | $3.49 \times 10^{-5}$ | 0.0001047 |
| **Copanlisib** | $1.09 \times 10^{-8}$ | $2.46 \times 10^{-8}$ | $3.28 \times 10^{-8}$ | $9.86 \times 10^{-8}$ |
| **Buparlisib** | $1.47 \times 10^{-7}$ | $3.29 \times 10^{-7}$ | $4.39 \times 10^{-7}$ | $1.32 \times 10^{-6}$ |
| **PI-103** | $4.94 \times 10^{-8}$ | $1.11 \times 10^{-7}$ | $1.48 \times 10^{-7}$ | $4.44 \times 10^{-7}$ |

**Supplementary figure 3. Kras G12A NSCLC-PDTOs sensitivity to target inhibitors.**
PDTOs were digested into a single cell suspension and cells were plated in a 384 well plate at a density of 1000 cells per well in 8ul droplets (1:2 media:matrigel). Plates were centrifuged briefly to ensure the cells were at the bottom of the well and 15µl of media were added. Cells were incubated for 72h to allow the cells to form PDTOs and afterwards the drugs were added and incubated for additional 96h. Concentrations ranging between 10 and 0 (serial 1/3 dilutions) were assessed for 23 different inhibitors against: EGFR (Lapatinib, Osimertinib, Dacomitinib, Afatinib, Erlotinib and Gefitinib), PI3K (Idelalisib, Copanlisib, PI-103, Buparlisib, GSK2636771, Parsaclisib and Erganelisib), mTOR (Temsirolimus, Rapamycin and Everolimus), ERK (Ulixertinib), Raf (AZ628 and Dabrafenib), Ras (AMF510) and MEK (Binimetinib,Trametinib and Selumetinib). The readout was performed using CellTiterGlo®3D reagent according to the manufacturer's protocol. Luminescence was measured by the Biotek Synergy H4 plate reader. Full dose-response curves are depicted for each patient in **A**. Each color represent a particular target. Dose response curves for the hit compounds and their estimated effective concentrations 10, 20,25 and 50 are depicted for WCM3409 (**B**) and for WCM3407 (**C**).

.

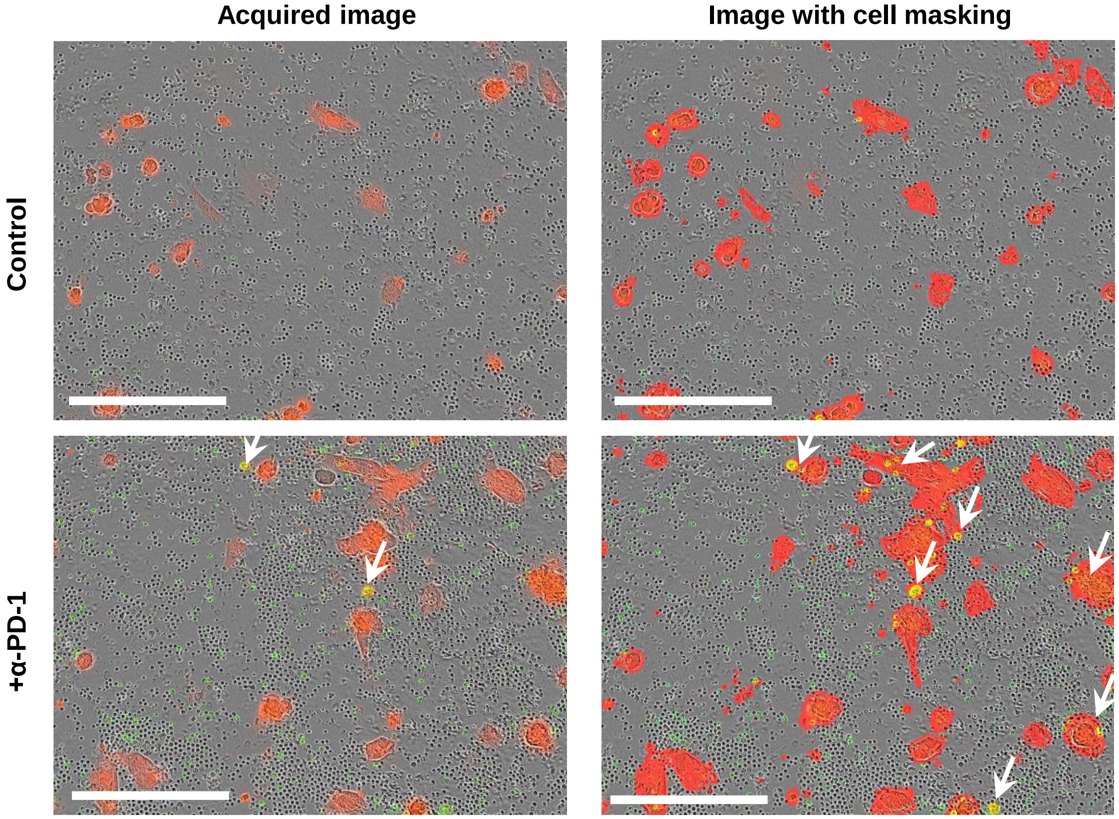


**Supplementary Figure 4. Representative cell masking setting for the identification of apoptotic PDTOs.** Single tumor cells suspensions were stained with cell far red following the manufacturer protocol and seeded in Matrigel 66% droplets. After 3 days of culture PDTOs were harvested using cell recovery solution and seeded in 96 wells plates were T-cells were added (3:1, Effector:Target) in T-cell media containing 5uM NucView488 caspase-3 substrate. Shown are representative images of killing assays with or without the addion of α-PD-1. On the left images are depicted as they are acquired by the Incucyte S3.On the right images are display with the corresponding cell masks for red events (PDTOs), green events (apoptotic T-cells) and yellow events (apoptotic PDTOs). Arrow heads indicate apoptotic PDTOs. Cell masking improve the visualization of the apoptotic PDTOS. Estimation of the percentage of apoptotic PDTOs in each experimental condition is determined normalizing the apoptotic PDTOs (yellow events)/field to the total PDTOs (red events)/field. Scale bar indicate 200µm.
