## Supplementary Table 1 for "Novel co-culture strategies of tumor organoids with autologous T-cells reveal clinically relevant combinations of immune-checkpoint and targeted therapies"

| Patient | Gender | Race/ Ethnicity | Smoking status | Cancer stage | Lesion size on pathologicic examination (cm) | T stage | N stage | M stage |
| --- | --- | --- | --- | --- | --- | --- | --- | --- |
| WCM3407 | M | White | Former | IIB | 4.0 | T3 | N0 | M0 |
| WCM2499 | F | White | Former | IIIA | 7,6 | T4 | N0 | M0 |
| WCM3606 | F | African American | Former | IIA | 3,8 | T2a | N1 | M0 |
| WCM3417 | F | White | Never | IA3 | 3 | T1c | N0 | M0 |
| WCM3409 | F | African American | Never | IA3 | 2,9 | T1c | N0 | M0 |
| WCM3410 | F | White | Former | IIIA | 7,3 | T4 | N0 | M0 |
| WCM3602 | M | White | Current | IIIA | 3,3 | T2a | N2 | M0 |
| WCM3603 | F | White | Never | IA2 | 1,7 | T1b | N0 | M0 |
| WCM3604 | M | White | Never | IB | 3,3 | T2a | N0 | M0 |
| WCM3605 | F | White | Never | IA3 | 2,6 | T1c | N0 | M0 |
| WCM3413 | M | White | Former | IIIA | 3 | T1b | N0 | M0 |
| WCM3066 | M | White | Current | IA2 | 1,9 | T3 | N1 | M1a |
| WCM3607 | F | White | Never | IA3 | 2,1 | T1c | N0 | M0 |
| WCM3416 | M | White | Former | IIB | 5,5 | T3 | N0 | M0 |
| WCM3608 | M | Asian | Never | IB | 3,5 | T2a | N0 | M0 |
| WCM3083 | F | unknown | Current | IVA | 4 | T3 | N1 | M1a |
| WCM3289 | M | Asian | Former | IIIB | 6,2 | T3 | N2 | M0 |

**Supplementary table 1. Patients clinical information.**
